# Choices Change the Temporal Weighting of Decision Evidence

**DOI:** 10.1101/2020.03.06.979690

**Authors:** Bharath Chandra Talluri, Anne E. Urai, Zohar Z. Bronfman, Noam Brezis, Konstantinos Tsetsos, Marius Usher, Tobias H. Donner

## Abstract

Decisions do not occur in isolation, but are embedded in sequences of other decisions, often pertaining to the same source of evidence. Here, we characterized the impact of intermittent choices on the accumulation of a protracted stream of decision-relevant evidence towards a final decision. Human participants performed two versions, based on perceptual or numerical evidence, of a decision task that required two successive judgments at different times during the evidence stream: an intermittent response consisting of a binary choice, and a continuous estimation at the end of the evidence stream. In a control condition, subjects executed a choice-independent motor response instead of binary choice as the intermittent response. In both, perceptual and numerical tasks, the intermittent choice reduced the sensitivity of subsequent evidence, and flipped the relative temporal weighting of early and late evidence in the final estimation judgment. The individual extent of the choice-induced overall (non-selective) sensitivity reduction predicted the extent of the selective down-weighting of subsequent evidence inconsistent with the initial choice, a form of confirmation bias. In sum, active decisions about a protracted evidence stream profoundly alter the dynamics of evidence accumulation, consistent with an active, modulatory mechanism triggered by the choice.

## INTRODUCTION

Many decisions are made under uncertainty, on the basis of noisy, incomplete, or ambiguous decision-relevant ‘evidence’. An extensive body of research on perceptual decisions under uncertainty has converged on the idea that evidence about the state of the sensory environment is continuously accumulated across time [1,2]. In the perceptual choice tasks commonly used in the laboratory (but see [3–5]), performance is maximized by weighing evidence equally across time [1]. Yet, the evidence weighting applied by human and non-human decision-makers often deviates substantially from such flat weighting profiles (but see [6,7]): some studies found stronger weighting of early evidence (‘primacy’; [8–11]), others stronger weighting of late evidence (‘recency’; [12–14]), and yet others even non-monotonic weighting profiles [15].

Most of these studies of perceptual choice have focused on the within-trial factors governing decision-making, ignoring interactions between consecutive decisions or stimuli. However, real-life decisions are not isolated events, but embedded in a sequence of judgments based on continuous streams of information. Indeed, a growing body of evidence has shown that perceptual choices are biased by the choices made on previous trials [16–32]. Recently developed task protocols provide new tools for assessing the impact of choices on the accumulation of subsequent decision evidence. These tasks prompt two successive judgments within the same trial: commonly a binary choice followed up by a continuous estimation [33–38] or a confidence [39,40] judgment. Specifically, some tasks prompt binary choice and estimation judgments sequentially, separated by a second evidence stream presented in between [35,39,40,38]. These task designs have led us to two insights. First, the overall sensitivity to evidence following the intermittent choice is reduced in a non-selective (‘global’) fashion, a finding obtained in the domain of numerical decisions [35]. Second, sensitivity for information consistent with the binary choice is selectively enhanced, at the expense of less sensitivity for choice-inconsistent evidence, a choice-induced evidence re-weighting that produced a bias to confirm the initial choice and that was found for both perceptual and numerical decisions [38]. In the latter study, we did not examine the non-selective impact of choice in reducing sensitivity for subsequent evidence.

Here, we re-analyzed the datasets from both the previous studies [35,38] to develop a more comprehensive understanding of these two choice-dependent effects (selective evidence re-weighting and non-selective sensitivity reduction), as well as their relationship. We tested for the following three outstanding issues: (i) if the choice-induced sensitivity reduction observed in the domain of numerical decisions generalizes to the domain of perceptual decisions; (ii) how an intermittent overt choice affects the *temporal weighting* of evidence, and (iii) if and how the overt choice-induced, overall reduction of sensitivity relates to the choice-induced confirmation bias towards consistent evidence.

## MATERIALS AND METHODS

### Perceptual task

Participants were presented with two random dot motion stimuli in succession, and were asked to estimate the average motion direction across the two intervals in each trial (Figure 1A). A white line plotted on top of the circular aperture served as the reference, whose position changed between trials. An auditory cue after the first interval instructed the participants to either (i) report a binary choice about the direction of dots in the first interval (clockwise or counter-clockwise w.r.t the reference; two-third proportion of all trials), or (ii) make a choice-independent button press (one-third proportion of all trials). This intermittent response allowed us to investigate if participants showed different sensitivity to the second stimulus depending on whether they reported a binary choice (so called “Choice trials”), or made a choice-independent motor response (so called “No-Choice trials”). The delay between the first and second stimuli was fixed (2 seconds), regardless of the reaction time of the subject. Half of all choice trials ended with an auditory feedback about the correctness of the binary choice to motivate participants to take the binary choice component seriously. The coherence of the stimuli was fixed at a pre-determined level for each subject, while the direction of coherent dots in the two intervals was sampled independently from five possible values (−20°, −10°, 0°, 10°, 20° relative to the reference line). 23 possible combinations of directions were used in the experiment (excluding the two most obviously conflicting directions: −20°/20° and 20°/−20°).

**Figure 1.**
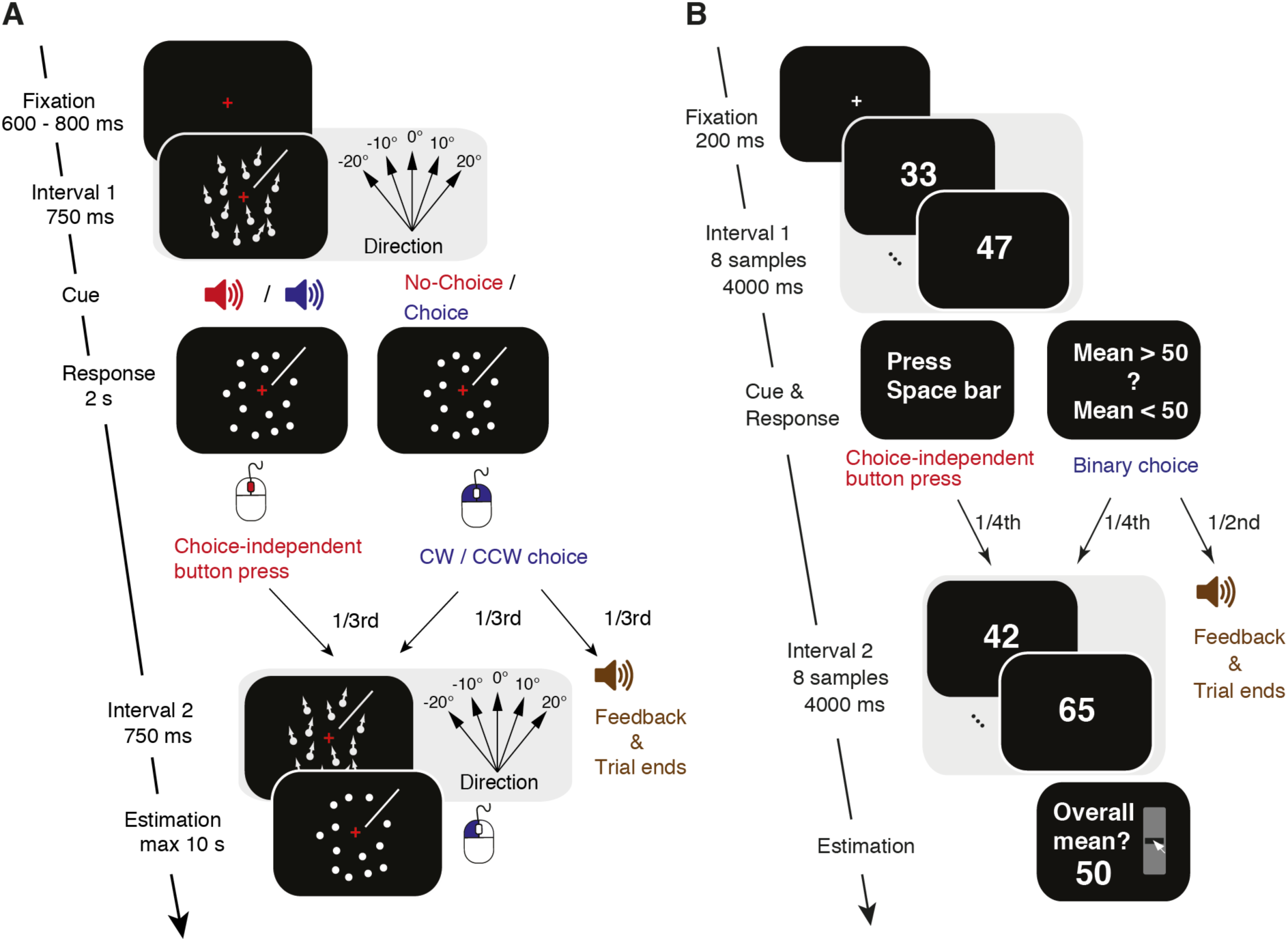
Behavioral tasks: Schematic of sequence of events within a trial. **A.** Perceptual task. After a first random dot motion stimulus was shown for 750 ms, participants received an auditory cue about whether to report a binary choice about the net motion direction (Choice trials) or to continue the trial (No-Choice trials). The choice entailed discriminating the motion direction as CW or CCW with respect to the reference (white line shown at about 45° in this example). On half the Choice-trials, auditory feedback was then given and the trial terminated. In the other half, and in all No-Choice trials, a second motion stimulus was presented (with equal coherence as the first, but independent direction), and participants were asked to report a continuous estimate of the mean direction of both stimuli by dragging a line along the screen with the mouse. See <u>here</u> for a video of the task structure. **B.** Numerical task. After the first sequence of eight numerical samples, participants were instructed to either press the space bar (a quarter of all trials; No-Choice trials), or to give their binary choice about the average of the eight samples (mean > or < 50; Choice trials). On two-thirds of Choice trials (constituting a half of all trials), auditory feedback was presented and the trial terminated. In the rest, a second sequence of eight numerical samples was presented and participants were instructed to give a continuous estimate of the mean across the two sequences. Adapted from [38] under a CC-BY license.

In all the analyses that follow, we used trials where participants made an estimation judgment (Choice trials and No-Choice trials). We excluded trials in which participants did not comply with the instructions i.e., when they pressed the mouse wheel on Choice trials or a choice key on No-Choice trials, trials in which the binary choice response time was below 200 ms, and trials where estimations were outliers. Outliers were defined as those trials whose estimations fall beyond 1.5 times the interquartile range above the upper-quartile or below the lower-quartile of estimations. Together, these excluded trials corresponded to ∼7% of the total trials across participants.

We analyzed data from the same set of participants as in our earlier report [38]. Please refer to this report for a detailed description of the task, participants, and stimuli used in the experiments.

### Numerical task

We reanalyzed data from the numerical integration task in [35] using the same analyses methods as the perceptual task. The task has a similar structure as the perceptual task above, with the exception that participants saw 16 numerical samples displayed in succession and reported their mean as a continuous estimate. Like the perceptual task, participants received a cue midway through the trial (i.e., after the first 8 samples) to either report a binary choice about the mean of the 8 samples (greater, or less than 50), or make a choice-independent motor response. In 50% of all trials, the trial terminated with feedback after the binary choice. On another 25% of the trials, participants saw the second stream of 8 numerical samples and made the continuous estimation judgment at the end (Choice trials). In the rest 25% of trials, participants made a choice-independent motor response (No-Choice trials) instead of the binary choice, and a continuous estimation judgment at the end. We analyzed data from all the trials where participants made the estimation judgment (50% of all trials).

We analyzed data from 20 out of 21 participants participated in the study. The remaining subject (subject 21) failed to do the task (Spearman’s correlation between estimation and mean evidence in No-Choice trials: rho = 0.18, p = 0.117; and in Choice trials: rho = 0.17, p = 0.156). Please see the earlier reports [35,38] for more detailed description of the task, stimuli, and participants.

### Pupillometry

Horizontal and vertical gaze position as well as pupil diameter were recorded at 1000 Hz using an EyeLink 1000 (SR Research). The eye tracker was calibrated before each block. Blinks detected by the EyeLink software were linearly interpolated from −150 ms to 150 ms around the detected velocity change. All further data analysis was done using FieldTrip [41] and custom Matlab scripts. We estimated the effect of blinks and saccades on the pupil response through deconvolution, and removed these responses from the data using linear regression. The pupil signal was bandpass filtered from 0.01 to 10 Hz using a second-order Butterworth filter, z-scored per block of trials, and down-sampled to 20 Hz. For both experimental conditions (Choice, No-Choice), we then averaged the time courses across trials, time-locked to either the onset of the evidence sequence.

### Modelling estimation reports

#### General approach

We used a statistical modelling approach to estimate the relative contributions of the sensory evidence (i.e., physical stimulus corrupted by sensory noise) conveyed by both successive dot motion stimuli to participants’ trial-by-trial estimation reports, as in our previous report [38]. The noisy sensory evidence was described by: 

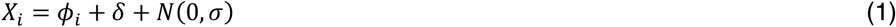

where *i* ∈ (1,2) denotes the interval, *ϕ*_*i*_ is the physical stimulus direction, *N*(0, *σ*) was zero mean Gaussian noise with variance *σ, δ* and *σ* were each observer’s individual overall bias and sensory noise parameters taken from the psychometric function fit to the binary choice data (see STAR methods in [38]).

### Global Gain model

We modelled a global, choice related reduction in sensitivity to evidence following an overt choice, by allowing the weights to vary separately in Choice trials and No-Choice trials. The estimations were modelled by:

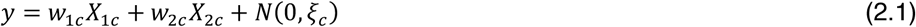

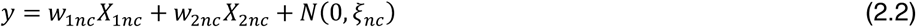

where *y* was the vector of estimations, *w*_1*c*_ (*w*_1*nc*_) and *w*_2*c*_ (*w*_2*nc*_) were the weights for the noisy evidence encoded in intervals 1 and 2 in Choice (No-Choice) trials respectively. *N*(0, *ξ*) was zero-mean Gaussian estimation noise with variance *ξ* that captured additional noise in the estimations, over and above the sensory noise corrupting binary choice.

### Fitting procedure

We used maximum likelihood estimates to estimate parameters and the goodness of fit for different models. To obtain the best fitting parameters that maximize the likelihood function of each model, we used the Subplex algorithm [42,43], a generalization of the Nelder-Mead simplex method, which is well suited to optimize high dimensional noisy objective functions. Please refer to our earlier report for a detailed description of the fitting procedure [38].

### ROC analysis for differences in sensitivity to evidence in interval 2

We assessed the impact of an overt choice on sensory evidence in interval 2 from participants’ estimations in a model-free fashion, using the so-called ROC analysis. This analysis was based on the receiver operating characteristic [44], similar to the one used in our earlier report (see “model-free analysis of estimation reports” in [38]). By computing ROC indices between sets of trials that differed in their input, we could assess the sensitivity of the observer in using that input to guide their estimation reports.

For the perceptual task, in each condition (Choice and No-Choice), we first sorted trials based on the directional evidence in interval 1 (*ϕ*_1_). For each *ϕ*_1_, we ran the ROC analysis on all pairs of estimation distributions, separated by 10° of directional evidence in interval 2 (*ϕ*_2_): −20° vs. −10°, −10° vs. 0°, 0° vs. 10°, and 10° vs. 20°. This gave us 4 ROC-indices per *ϕ*_1_, one index for every pair of distributions compared. We then computed a weighted average ROC-index for each *ϕ*_1_, weighting the individual ROC-indices by the number of trials that went into the ROC analysis. The resulting ROC indices, which are robust to changes in *ϕ*_1_, are averaged to obtain one single ROC index per subject for each condition.

ROC indices for the numerical task were computed similar to the above procedure with the following exceptions: mean evidence in interval 1, and interval 2 were binarized (mean > 50 or mean < 50) resulting in two bins for interval 1, and interval 2 respectively. These binarized values were treated equivalent to *ϕ*_1_ and *ϕ*_2_ in the perceptual task above.

### Statistical tests

Non-parametric permutation tests [45] were used to test for group-level significance of individual measures for each task, unless otherwise specified. This was done by randomly switching the labels of individual observations either between two paired sets of values, or between one set of values and zero. After repeating this procedure 100,000 times, we computed the difference between the two group means on each permutation and obtained the p value as the fraction of permutations that exceeded the observed difference between the means. All p values reported were computed using two-sided tests, unless otherwise specified.

To obtain the correlation values for data pooled from both the tasks (Figure 4, 5), we first obtained Pearson’s correlation coefficient for dataset from each task (also reported in the figures). We then applied Fisher-transformation on the correlation values, calculated their weighted average to obtain the pooled Fisher-transformed correlation coefficient. This quantity is used to obtain the pooled Pearson’s correlation coefficient (using inverse Fisher transformation), and its corresponding p-value.

**Figure 2.**
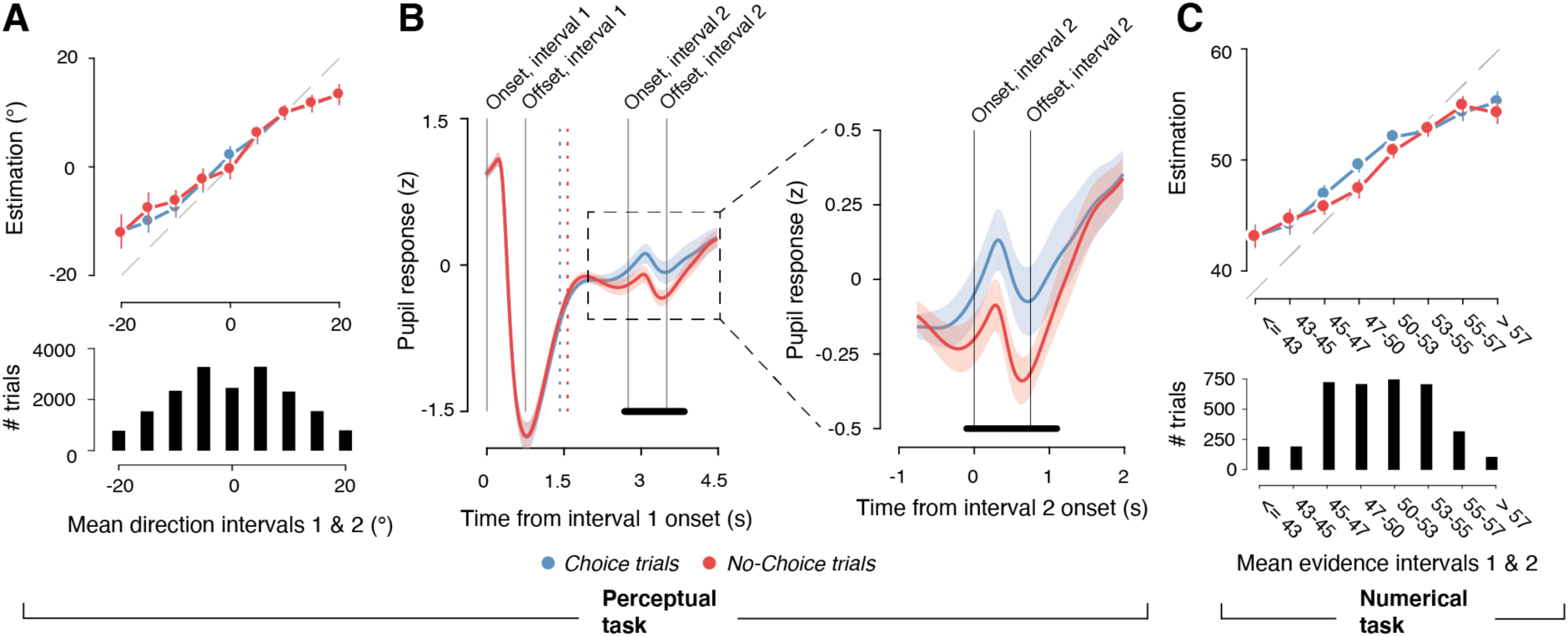
Behavior in both tasks. **A, B.** Perceptual task. **C.** Numerical task. **A.** Top: Continuous estimations as a function of mean direction across both stimuli. Bottom: Distribution of mean directions across trials. Data points, group mean; error bars, SEM. Stimulus directions and estimations were always expressed as the angular distance from the reference, the position of which varied from trial to trial (0° equals reference). **B.** Time courses of average pupil diameter aligned to trial onset for Choice and No-Choice conditions in Perceptual task. Left, average time course across whole trial. Right, close-up of time course during second stimulus interval (following intermediate motor response). Dashed vertical lines, mean response times across participants; grey vertical lines, different events during the trial. **C.** Same as A but for Numerical task. Mean evidence across intervals 1& 2 in C split into discrete bins for illustration. All panels: solid lines, mean across participants; shaded region, SEM; black horizontal bars, p<0.05, cluster-based permutation test Choice vs. No-Choice.

**Figure 3.**
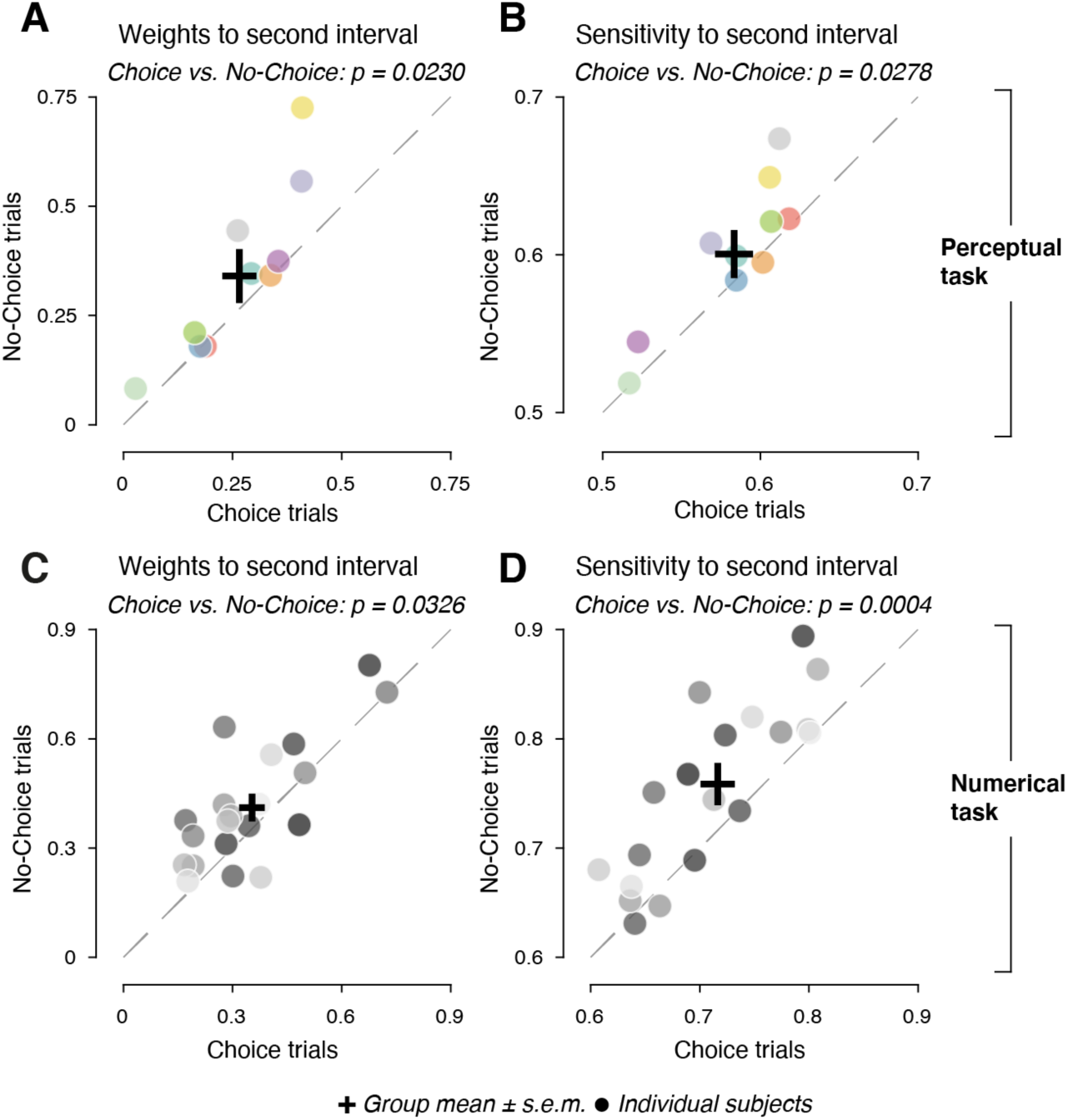
Sensitivity to second stimuli in Choice and No-Choice trials. **(A)** Model weights for sensitivity to second interval in Choice and No-Choice conditions in Perceptual task. Dashed line, identity of Choice and No-Choice; points above diagonal indicate larger weights to No-Choice. **(B)** As (A), but for ROC indices quantifying the sensitivity to second interval in a model-free way in Perceptual task. Dashed line, identity of Choice and No-Choice; points above diagonal indicate greater sensitivity to No-Choice. Data points, individual participants, with identical color scheme from (A, B). **(C, D)** As (A, B), but for Numerical task. Perceptual task, n = 10 participants; Numerical task, n = 20 participants; p values, permutation tests across participants (100,000 permutations).

**Figure 4.**
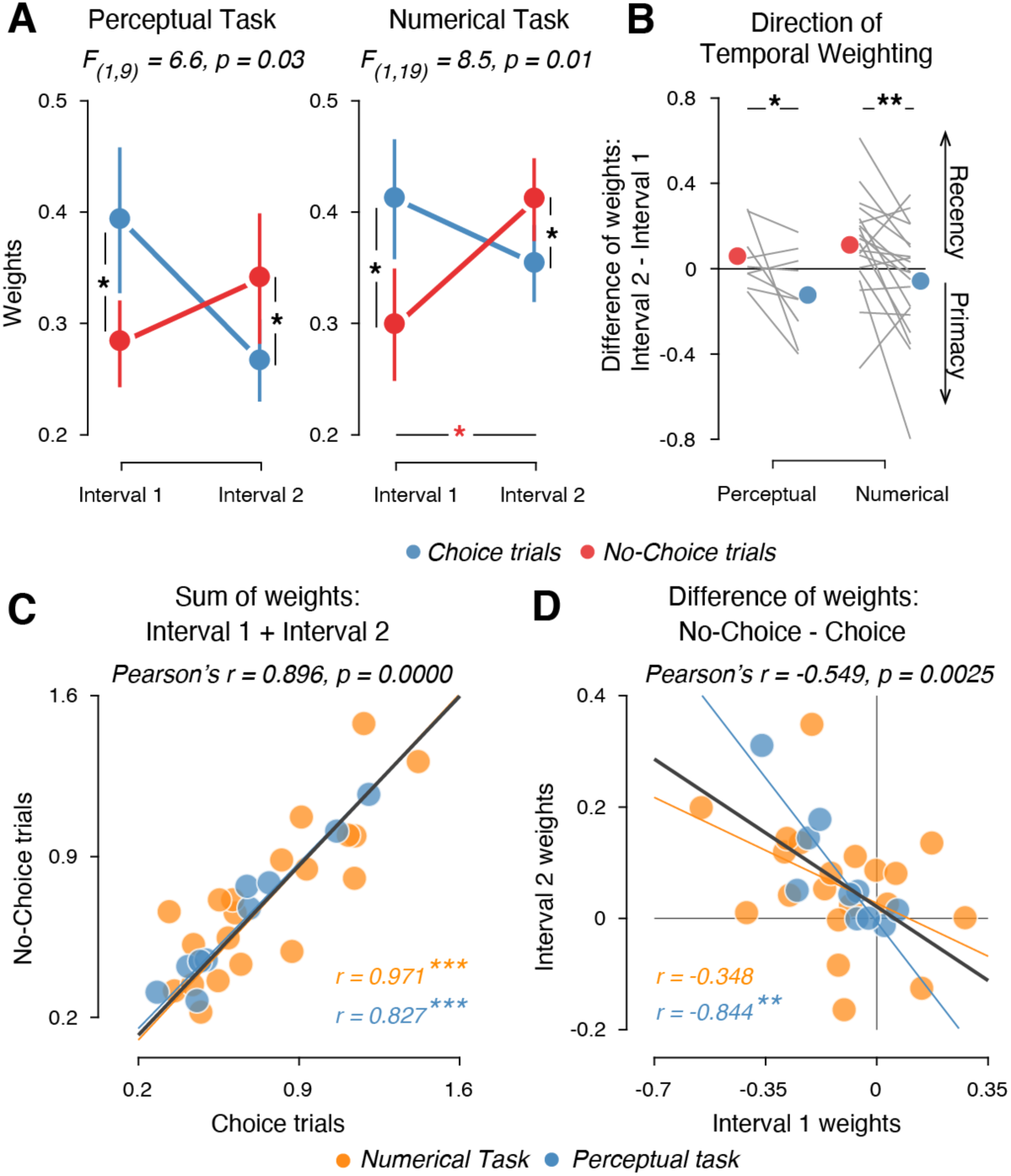
Choice-dependent alteration of temporal weighting profiles. **A.** Mean model weights for both stimulus intervals in Choice and No-Choice conditions in Perceptual task (left, n = 10 participants) and Numerical task (right, n = 20 participants). Error bars, SEM; F-statistic, interaction between interval and condition (repeated measures 2-way ANOVA). **B.** Direction of temporal weighting quantified as difference in model weights between interval 2 and interval 1, separately for each task; * p < 0.05, ** p < 0.005, permutation tests across participants (100,000 permutations). **C.** Sum of weights across both intervals in Choice and No-Choice conditions, across participants from both tasks. **D.** Difference between weights in Choice condition and No-Choice condition, in both intervals across participants from both tasks. Data points, individual participants; solid lines, best fitting lines; r, Pearson’s correlation coefficients; * p < 0.05, ** p < 0.005, *** p < 0.0005.

**Figure 5.**
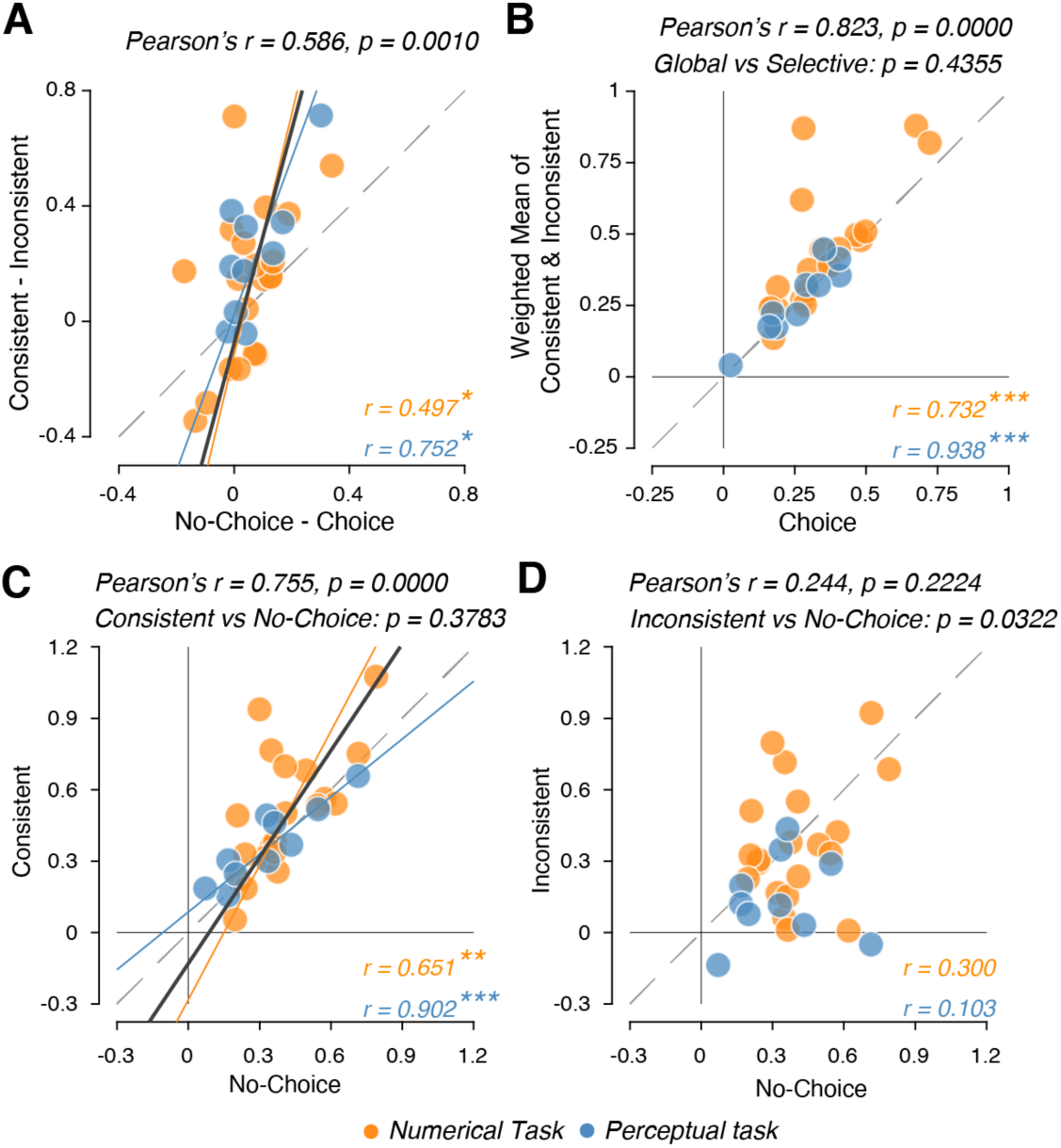
Relationship between global and consistency-dependent sensitivity modulation. **A.** Relationship between global (quantified as the difference in weights to second interval between No-Choice and Choice conditions) and selective sensitivity modulation giving rise to Confirmation bias (quantified as the difference in weights to second interval between Consistent and Inconsistent conditions, from [38]), across participants from both tasks. **B.** Relationship between the weighted mean of Consistent and Inconsistent conditions (from [38], weighted by the number of trials in each condition) and weights for the Choice condition for the second interval across participants from both tasks. **C, D.** Relationship between No-Choice weights for the 2nd interval; and Consistent weights (C), and Inconsistent weights (D). Data points, individual participants; solid lines, best fitting lines; dashed lines, identity lines; p-values comparing the two conditions in B, C & D are obtained from 2-way ANOVA with the conditions and tasks as factors; * p < 0.05, ** p < 0.005, *** p < 0.0005.

## RESULTS

Participants reported a continuous estimate of the mean of fluctuating sensory (perceptual task, Figure 1A) or symbolic (numerical task, Figure 1B) evidence across two successive intervals. This estimate needed to be based on accumulating some internal representation of the fluctuating evidence – motion direction or numerical value in the perceptual or numerical tasks, respectively – across the two stimulus intervals. On a subset of trials (so-called Choice trials), participants were also asked to report an intermediate categorical choice after the first stimulus: a fine direction discrimination judgment relative to a visually presented reference line (Perceptual Task) or comparison of the numerical mean with 50 (Numerical Task). On the remaining set of trials (No-Choice trials), participants were asked to simply press a button for continuing the trial, without reporting a categorical judgment of the first evidence. The cue informing participants whether to report the discrimination judgment or to press a choice-independent button press came after the first stimulus interval. This design enabled us to quantify the degree to which evidence in each interval contributed to the final estimation and whether this depended on the overt report of a categorical choice (Materials and Methods).

Estimation responses in both tasks increased with mean directional evidence across the two intervals (Figure 2A, 2C), and did not differ between Choice and No-Choice trials, with negligible and statistically non-significant differences in the regression slopes for evidence against estimations (perceptual task: 0.0256, p = 0.8449; numerical task: 0.0125, p = 0.9083). Participants’ pupils constricted after trial onset in the same way for Choice and No-Choice trials, an expected response to the onset of the random dot stimulus (pupil light reflex, Figure 2B; pupil diameter was only measured during the perceptual task). This constriction was followed by a dilation about 1 s after the intermittent response (see red/blue dashed vertical lines), indicating a phasic activation of central arousal systems [46–49]. Critically, this dilation was bigger for Choice than No-Choice trials (Figure 2B), reflecting the internal decision process associated with Choice [47,50]. Indeed, the bigger pupil dilation during Choice was not due to longer response times in that condition (and the associated longer accumulation of central inputs in the peripheral pupil apparatus; [47,50]): response times were, in fact, shorter in Choice than No-choice trials (see blue and red vertical lines in Figure 2B; permutation test, p = 0.0112).

### Global down-weighting of subsequent evidence following intermittent choice

We first replicated our finding, previously reported for the Numerical Task [35], of lower sensitivity to subsequent evidence in the Choice condition, for the Perceptual Task (Figure 3). A statistical model-based as well as a model-free approach (Materials & Methods) both showed a choice-dependent sensitivity reduction for subsequent evidence (Figure 3). Model weights for the second stimulus were significantly smaller in Choice trials compared to No-Choice trials (most individual participants, and the mean, above identity line in Figure 3A, 3C). Likewise, a model-free measure of sensitivity to subsequent evidence (area under the ROC curve) was smaller on Choice trials compared to No-choice trials (most individual participants, and the mean, above identity line in Figure 3B, 3D). In sum, overt choices reduce the sensitivity to subsequent evidence not only for numerical, but also for perceptual decisions.

### Intermittent choice alters temporal weighting of sensory evidence

Having generalized the choice-induced sensitivity reduction across both domains of decision-making, we assessed if and how the intermittent choice affected the relative weighting of early vs. late evidence in the decision process underlying the final estimation judgments. For both tasks, the weights in Choice trials were higher for the first interval, and lower for the second interval compared to No-Choice trials, with a significant interaction between trial type (Choice vs. No-Choice) and interval (Figure 4A). In other words, the evidence weighting across the two intervals flipped from recency to primacy between No-Choice and Choice conditions (Figure 4B). This choice-induced flip in temporal weighting was also evident in the individual data: The sums of weights from both intervals were highly similar for Choice and No-Choice trials in each subject (Figure 4C), but the *difference* in Choice and No-Choice weights was negatively correlated between intervals (Figure 4D). Please note that no such constraint was imposed in the statistical models used to estimate the weights for both intervals (Materials and Methods).

These results are in line with a ‘push-pull’ mechanism, in which a limited resource was distributed across sensitivity to evidence in both intervals: Reporting an intermittent choice after the initial evidence boosted sensitivity to that early evidence, but at the cost of reducing sensitivity to subsequent evidence. This effect could also explain the similarity in overall estimation accuracy between Choice and No-Choice conditions (Figure 2A, 2C).

### Choice-dependent non-selective, and selective sensitivity modulations are coupled

Finally, we found that the individual degree of the choice-dependent, overall (non-selective) reduction in sensitivity to subsequent evidence was closely related to the selective confirmation bias effect defined as a larger sensitivity to subsequent evidence consistent than inconsistent with the initial choice (Figure 5). We quantified the overall (‘global’) gain modulation as the difference in weights of interval 2 between Choice trials and No-Choice trials, and the selective choice-driven gain modulation as the difference in weights of interval 2 between trials with choice-consistent and -inconsistent evidence. Participants with a stronger global gain reduction also showed a stronger selective gain modulation (Figure 5A). The weights for interval 2 in the Choice condition in Figure 3A, 3C are indeed the weighted average of the corresponding weights in choice-consistent and -inconsistent evidence (Figure 5B). Furthermore, we found that the weights for interval 2 in the No-Choice condition were comparable in magnitude, and correlated across participants with the weights for choice-consistent evidence (Figure 5C), but not for choice-inconsistent evidence (Figure 5D). Thus, the reduction in sensitivity following an overt choice observed in Figure 3 was primarily driven by the reduction in sensitivity to evidence inconsistent with the initial choice.

## DISCUSSION

Recent work has begun to expose the impact of choices on the accumulation of subsequent decision evidence, revealing an overall reduction in sensitivity to subsequent evidence [35] combined with a selective suppression of the gain of evidence inconsistent with the initial choice (confirmation bias) [38]. Here, we extend this nascent line of work, by showing that the report of an intermittent choice about a protracted stream of perceptual or symbolic (numerical) evidence alters the weights assigned to pre- and post-choice evidence from recency to primacy. Further, we show that the above three effects are tightly related, consistent with generation by the same mechanism.

The overall reduction in sensitivity due to the initial choice observed here (Figure 3) corroborates earlier analyses of the Numerical Task data [35]. The correspondence between these and our current findings from the Perceptual Task indicate that choice-induced decreases in sensitivity generalizes across different formats of decision evidence (from symbolic to low-level perceptual). We used the same methods to analyze data from the perceptual task presented here (but different from those in [35], see Materials and Methods), and data from the numerical integration task using a similar task protocol [35], and found strong correspondence between the two.

One important implication of our findings is that the temporal weighting profiles in evidence accumulation are neither fixed traits of decision-makers nor fixed properties of certain tasks, but are flexibly altered on the fly in a given task, depending on the presence or absence of an intermittent choice. Previous studies investigating temporal biases in decision-making found conflicting results, ranging from recency to flat profiles, to primacy. Differences in the task protocols and idiosyncratic tendencies of decision-makers are important confounds that complicate the comparison between these studies. Our current results show that the temporal weighting profile, within a given individual and a given task, can be effectively flipped, simply by asking the participant for an intermittent choice half-way through the evidence stream.

Importantly, the here-discovered, strong relationship between the individual strength of the choice-induced, global sensitivity reduction and choice-induced, selective confirmation bias (Figure 5) is not a given, because both effects were operationalized in terms of two orthogonal comparisons: the sensitivity reduction by comparing sensitivity between trials with an intermittent choice and trials without such a choice; the confirmation bias by comparing trials with subsequent evidence that was consistent or inconsistent with the choice, *within* the trials that contain an intermittent choice. Thus, presence of a global sensitivity reduction effect does not imply presence of the confirmation bias, and vice versa. Even so, their correlations were tight, in line with a common underlying mechanism. In particular, the weights of new evidence in the No-Choice condition were comparable to, and correlated with the corresponding weights for the choice-consistent but not for choice-inconsistent evidence in the Choice-condition. This observation suggests a distinct state of the decision-maker when faced with information inconsistent with previous decisions, possibly reflecting the suppression of post-decisional dissonance [51].

It is tempting to interpret our findings as a signature of decision-related cortical evidence accumulation dynamics [52–54], combined with neuromodulatory input [5,47,55]. Once the decision circuits have settled in an attractor (choice commitment), this will reduce the decision-maker’s sensitivity to all subsequent evidence (see [35], Supplement) – an effect that may hold regardless of whether that evidence is consistent or inconsistent with the choice. Due to selective feedback from the accumulator circuit to early sensory regions encoding the evidence, the attractor state in accumulator networks may additionally cause selective gain modulation of subsequent incoming evidence [54] that can translate into the consistency-dependent, selective confirmation bias effect we found earlier [38]. These effects may have been amplified by choice-induced, phasic neuromodulatory input to cortex. Our task entailed an interrogation protocol, in which a categorical choice was prompted by the experimenter, when cortical decision circuits might not yet have reached a stable decision commitment; the choice prompt might then trigger the release of certain neuromodulators that pushes the decision circuits into an attractor state [1,35]. Such a neuromodulatory signal might be reflected in pupil dilation [47,48,56], which we found to be larger during the choice, compared to the no-choice condition. In sum, by altering the dynamical properties of decision circuits in the brain, choices can have versatile and coupled effects on evidence accumulation.

## ACKNOWLEDGEMENTS

We thank Ana Vojvodic for help with data collection. This research was supported by the German Academic Exchange Service (DAAD) (to AEU), a European Research Council Starting Grant under the European Union’s Horizon 2020 research and innovation program (Grant No. 802905) (to KT), and the following grants from the Deutsche Forschungsgemeinschaft (DFG, German Research Foundation): DO 1240/2-1 and DO 1240/3-1 (to THD). We acknowledge computing resources provided by NWO Physical Sciences.

## AUTHOR CONTRIBUTIONS

Conceptualization, T.H.D. and M.U.; Methodology, B.C.T., A.E.U., Z.Z.B., N.B. and K.T.; Software, B.C.T., A.E.U., and K.T.; Formal analysis, B.C.T., and A.E.U.; Investigation; B.C.T. and A.E.U.; Writing-original draft, B.C.T. and T.H.D.; Writing-review & editing, B.C.T., A.E.U., Z.Z.B., N.B., K.T., M.U., T.H.D.; Visualization, B.C.T.; Supervision, T.H.D., M.U. and K.T.; Project administration, T.H.D. and M.U.; Funding acquisition, T.H.D.

